# Cardiomyocytes execute pro- and anti-inflammatory signaling of IFNγ-induced GBP5 by differential regulation of the inflammasome

**DOI:** 10.64898/2026.03.16.712087

**Authors:** Laura Neuberger, Luisa Lange, Sandra Hoffmann, Timon Seeger, Lorenz Lehmann, Norbert Frey, Manju Kumari

## Abstract

Infiltration of conventional immune cells has been ascribed as the fundamental drivers of innate immune signaling in the damaged myocardium. However, the emerging intrinsic immunoregulatory potential of cardiomyocytes still remains poorly understood. Interferon gamma (IFNγ) is a pleiotropic cytokine with context-dependent detrimental as well protective role in regulating cardiac inflammatory circuits. The prevailing view of IFNγ as a prime pro-inflammatory cytokine has been challenged due to its paradoxical actions both as an inducer as well as negative regulator of inflammation, but the players involved in these converse processes remains enigmatic. Here we show that cardiomyocytes exhibit a cell-autonomous immunocompetent response upregulating innate inflammatory signaling upon type I and type II IFN stimulus. Notably, hiPSC-derived cardiomyocytes display a robust increase in guanylate binding protein 5 (GBP5), one of the major IFNγ-induced GTPase involved in inflammasome signaling, followed by upregulation of AIM2/CASP1 pathway whereas NLRP3 levels remain unaltered by IFNγ stimulation. GBP5 knockdown and overexpression studies in hiPSC-derived cardiomyocytes identify GBP5/TGFβ axis as a non-canonical anti-inflammatory feedback regulation on the IFNγ-induced inflammatory cascade.

## Introduction

Elevated IFNγ signaling plays an instrumental role in modulating innate immunity and renders cardiomyocytes vulnerable to a plethora of inflammatory responses in the injured myocardium^1, 2^. Our understanding of the inflammatory components regulating IFNγ response in the ischemic, non-ischemic, and anti-cancer drugs associated cardiomyopathies has emerged primarily from the studies elucidating function of immune cells, fibroblasts, or endothelial cells within myocardium whereas the intrinsic immunoregulatory potential of cardiomyocytes remains unclear.

IFNγ is the sole type II IFN and signals through heterodimeric IFNγ receptor to activate JAK/STAT1 pathway^3^. Nearly all cell surfaces express IFNγ receptor composed of two sub units, IFNGR1 and IFNGR2, and can thus respond to IFNγ stimuli. Lymphoid cells (T cells, natural killer T cells, and natural killer cells) and myeloid cells (CD11b+ macrophages) are the major source of IFNγ^3–5^. Of note, IFNγ levels are subject to intense transcriptional control by positive (T-bet, NFAT, Eomes, etc.) and negative (Prox1, TGFb, GATA3, etc.) transcriptional control factors to maintain cellular inflammatory homeostasis. Pro-inflammatory cytokines such as IL12, IL15 and IL18 function as the inducers of IFNγ expression whereas anti-inflammatory cytokines such as IL10 and TGFb limit the downstream inflammatory pathways by attenuating IFNγ production from T cells and natural killer cells.

Numerous research findings have established a central role for IFNγ as an inducer of multiple gene networks altering inflammation and inflammasome signaling in various cardiomyopathies. Studies from *Ifnγ*^-/-^ mice suggest that IFNγ exhibits protective role in sustained pressure overload induced cardiac hypertrophy^6, 7^. Similarly, IFNγ deficiency diminished cardiac myeloid cell influx, impaired cardiac function, and survival post myocardial infarction^8^. On the other hand angiotensin II-induced cardiac damage was ameliorated by IFNγ blockade^9^. Thus, IFNγ can augment or suppress inflammatory response in a context- and disease-specific manner. However, there is poor understanding of the players involved in this complex transition of context dependent paradoxical actions of IFNγ signaling as pro- vs anti-inflammatory cytokine^10, 11^.

Evidence for upregulation of inflammasome signaling in the pathophysiology of cardiac dysfunction has emerged from the research across numerous laboratories in the last decade^12, 13^. Inflammasomes are multimeric protein complexes that recognize various pathogen-associated molecular patterns (PAMPs) and damage-associated molecular patterns (DAMPS) and thus functions as internal sensors and receptors. Pattern recognition receptors (PRRs) such as Toll-like receptors (TLRs), and C-type lectin receptors respond to the extracellular space whereas NOD-like receptors (NLRs), absent in melanoma2 (AIM2)-like receptors, retinoic acid inducible gene-I (RIG-1)-like receptors recognize endogenous danger molecules^2^. Among these PRRs, a central role of *Nlrp3* upregulation during pressure overload caused by transverse aortic constriction (TAC) and in myocardial infarction model by left anterior coronary artery ligation (LAD) has been highlighted^13, 14^. Recently, AIM2 has also emerged as a critical inflammasome in mice and humans regulating atherosclerotic cardiovascular disease, ischemic and non-ischemic cardiomyopathy^15–17^.

IFNγ regulates inflammasome activation by primarily inducing GBPs expression as part of the interferon-stimulated gene (ISG) response. Among them, GBP5 belongs to a family of dynamine-related large GTPases that play an important role in canonical and non-canonical inflammasome activation in various infectious diseases, immune-checkpoint inhibitor induced myocarditis, cancers and immune disorders^18–20^. Importantly, GBP5 deficient mice corroborated defects in NLRP3 activation from bone-marrow derived macrophages only in response to PAMPs but not DAMPs suggesting differential pathway regulation at play upon sterile inflammation^21^. Divergent role of IFNγ and inflammasome (IL1β, NLRP3 and AIM2) regulation has led to a cognate quest for developing various therapeutic tools. Multiple clinical trials in developing inflammasome-targeting therapies are under evaluation targeting various cardiomyopathies (systolic heart failure, acute myocarditis, reperfusion injury, etc.)^22,23^. Therefore, it is imperative to identify the cardiomyocyte-specific unique and divergent regulatory pathways involved in exerting beneficial vs. detrimental effects of IFNγ-induced inflammasome signaling.

Here we show that cardiomyocytes execute the pro- and anti-inflammatory response of IFNγ in parallel to generate rheostat towards physiological inflammation. Cardiomyocytes exhibit differential regulation of inflammasome signaling and GBP5 expression towards type I and type II IFN activation. Importantly, our studies highlight the immuno-competence of cardiomyocytes by displaying a robust, cell-autonomous regulation of *GBP5/AIM2/CASP1* mediated inflammasome signaling in response to IFNγ whereas NLRP3 levels remain unaltered. Furthermore, GBP5 knockdown in hiPSC-derived cardiomyocytes suggest that IFNγ primed cardiomyocytes upregulate *TGFb* levels to combat aberrant activation of pro-inflammatory signaling and generate a controlled inflammasome response.

## Results

### Differential regulation of inflammasome signaling in neonatal rat cardiomyocytes by type I and type II IFN stimulation

Inflammasome signaling has emerged as a central regulator of cardiac damage response. Traditionally myeloid cells are known as effectors of inflammatory response. We and others have shown a key role of cardiomyocytes in regulating sterile inflammation^24–26^. However, a potential role of cardiomyocytes as an immunocompetent cell regulating inflammasome signaling upon IFN stimulation remains ill-defined. To investigate the cell-autonomous effect of elevated type I and type II IFN signaling on cardiomyocytes inflammasome activation, isolated cardiomyocytes from neonatal rats (NRCMs) were treated with interferon gamma (IFNγ) to stimulate type II IFN signalling and Poly: IC or lipopolysaccharide (LPS) to stimulate type I IFN pathway (Fig. 1a). Type II IFN stimulation by IFNγ showed robust upregulation of *Aim2*, *Casp1* and *IL1b* expression, however, failed to upregulate *Nlrp3* levels (Fig. 1b). On the other hand, type I IFN stimulation led to increased canonical inflammasome signaling with elevated expression of *Nlrp3*, *Casp1* and *IL1b* in cardiomyocytes (Fig. 1c-d). Upregulation of pro-inflammatory STAT1 signalling along with inflammasome pathway is a hallmark of interferon stimulation in immune cells. Our results show that type I and type II IFN priming led to increased pro-inflammatory state in primary cardiomyocytes with increased expression of *Stat1*, *Ccl2*, and *Ccl5* at mRNA (Fig. 1e-g). Importantly, this was associated with robust phosphorylation of STAT1 (Tyr701) at protein level (Fig. 1k-l). TGFβ and IL10 are anti-inflammatory cytokines that are often activated in parallel to an exaggerated inflammatory state in immune cells in order to counter balance and maintain cellular homeostasis^27^. However, these remain unaltered in IFN-stimulated neonatal rat cardiomyocytes suggesting a cell-type specific phenomena and a role for alternative pathways at play (Fig. S1a-S1c). Immune checkpoint inhibitor induced myocarditis and diabetic cardiomyopathy is associated with IFNγ-induced GBP5 and GBP6 expression in the myocardium as part of the interferon-stimulated gene (ISG) response^18, 28^. Here we show that cardiomyocytes with IFNγ stimulation elicited robust upregulation of *Gbp5* at mRNA and protein level compared to type I IFN stimulation (Fig. 1h-j). To note, IFNγ stimulation led to swift increase in GBP5 protein levels starting from 2h, wherein mRNA and protein levels reached peak after 6h and 12h of IFNγ stimulation, respectively (Fig S1d-S1f). However, both type I and type II showed modest stimulation of *Gbp6* compared to *Gbp5*. Additionally, due to reproducible robust effect on GBP5 and lack of specific antibody against GBP6 we focussed further studies with GBP5. Myocardial inflammation and heart failure are characterised by elevated inflammation and often associated with inflammasome activation in the failing hearts^2, 15^. Therefore, we tested whether GBP5 levels are altered in the failing heart with myocardial infarction (ischemic) and cardiac hypertrophy (non-ischemic). We found that mice subjected to ischemic cardiomyopathy by left anterior coronary artery ligation (LAD) for 4 days or cardiac hypertrophy by O-ring aortic banding (ORAB) for 2 weeks exhibited increased expression of GBP5 in the left ventricle tissue suggesting an increased inflammasome signaling (Fig. 1m-p). Taken together, these results suggest that cardiomyocytes from neonatal rats attain a robust, cell-autonomous, immunocompetent role in response to IFNγ by regulating GBP5/AIM2/CASP1 mediated inflammasome signaling.

**Figure 1.**
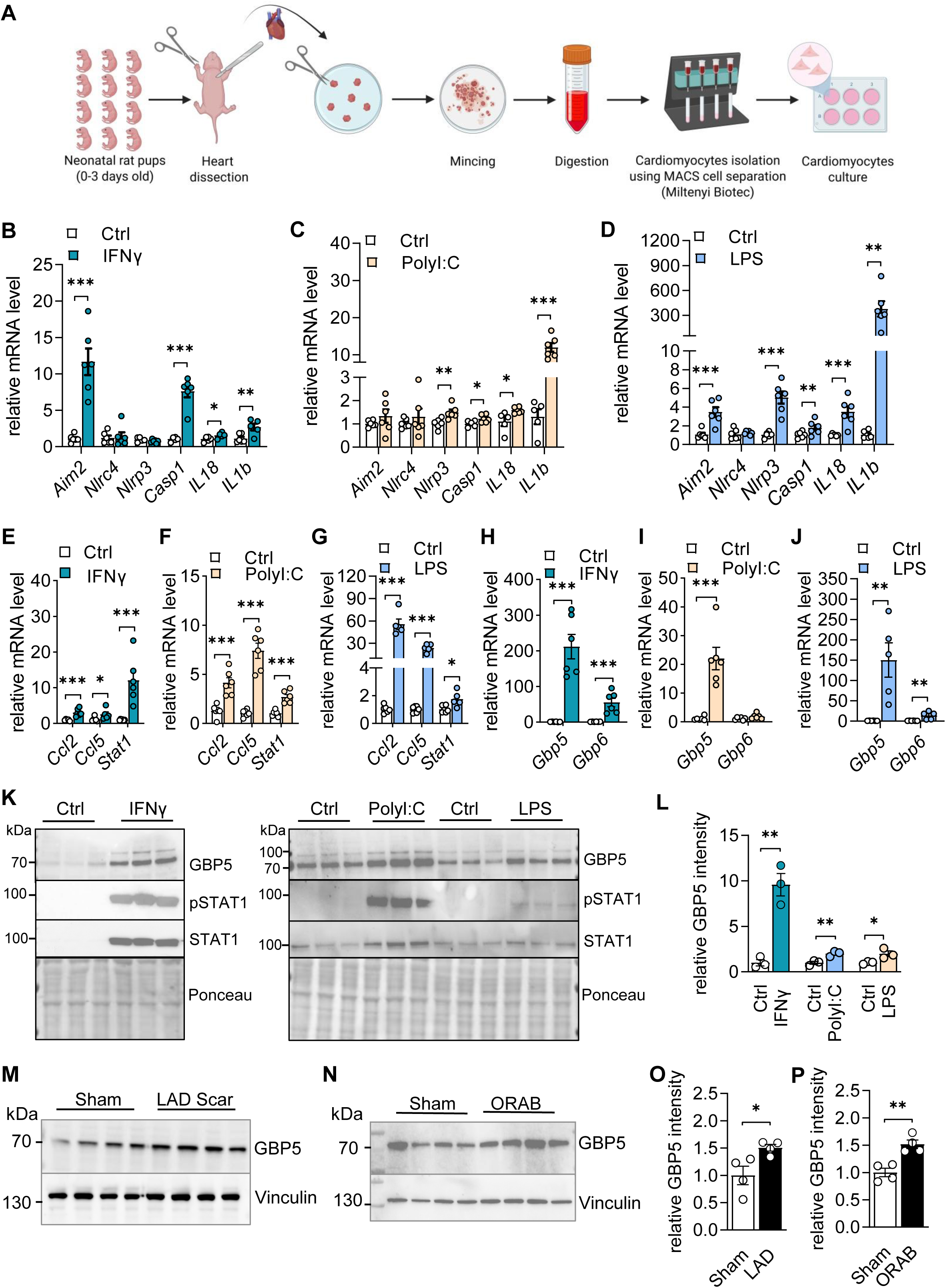
Cardiomyocytes exhibit robust upregulation of inflammasome signaling upon IFN activation. **A.** Schematic representation of isolation of neonatal rat ventricle cardiomyocytes (NRVCMs) following instructions of Miltenyi kit. Images prepared using BioRender (https://BioRender.com). **B-D.** Expression of inflammasome marker genes upon stimulation of NRVCMs with type I (LPS and PolyI:C) and type II (IFNγ) IFN stimuli. N=6/group. ***p<0.001, **p<0.01, *p<0.05. **E-G.** Expression of inflammatory marker genes upon stimulation of NRVCMs with type I (LPS and PolyI:C) and type II (IFNγ) IFN stimuli by qPCR. N=6/group. ***p<0.001, *p<0.05. **H-J.** Gene expression of *Gbp5* and *Gbp6* upon stimulation of NRVCMs with type I (LPS and PolyI:C) and type II (IFNγ) IFN stimuli by qPCR. N=6/group. ***p<0.001, **p<0.01. **K.** Immunoblot showing GBP5, pSTAT1, and STAT1 levels in NRVCM upon stimulation with type I (LPS and PolyI:C) and type II (IFNγ) IFN stimuli. **L.** Quantification of the GBP5 protein levels shown in the immunoblots of Figure 1K. N=3/group. **p<0.01, *p<0.05. **M.** Immunoblot showing GBP5 protein levels in left ventricle of mice after 4 days of ischemic cardiomyopathy induced by ligation of the left anterior descending coronary artery (LAD). **N.** Immunoblot showing GBP5 protein levels in the left ventricle of mice after two weeks of cardiac hypertrophy induced by 2 weeks of pressure overload utilizing ORAB method. **O-P.** Quantification of the immunoblots shown in Figure 1M and Figure 1N, respectively. N=4/group. **p<0.01, *p<0.05.

### Human iPSC-derived cardiomyocytes display early increase in CASP1 followed by GBP5/AIM2 expression upon IFNγ stimulation but fails to elevate NLRP3

Despite structural similarities, the IFNγ-receptor in rodents and humans lack species cross-reactivity and species-specific variation in the inflammasome components may further render context dependent alterations in IFNγ signaling within cardiomyocytes^29–31^. Additionally, differential expression of human and murine TLR transcripts in different cell types and variable transcript regulation plays a critical role in the cellular activation of inflammasome^32,33^. Of note, a comparative analysis determining effect of type I and type II IFN stimulation on inflammasome signaling in human induced pluripotent stem cell (hiPSC)-derived cardiomyocytes is not known. Here, we found that hiPSC-dervided cardiomyocytes show robust phosphorylation of STAT1 after 2 h of IFNγ priming (Fig. S2a). This was associated with increased expression of pro-inflammatory marker genes and a robust increase in *CASP1* expression (Fig. 2a, b). Despite early increase in STAT1 phosphorylation, unlike NRVCMs, hiPSC-derived cardiomyocytes showed a delayed response towards GBP5 protein expression with a peak in GBP5 mRNA levels at 24 h of IFN priming (Fig. 2c, d; Fig. S2a, S2b). Similar to NRVCMs, hiPSC-CMs fail to upregulate *NLRP3* upon IFNγ stimulation (Fig. 2a). Immunofluorescence study show that IFNγ led to nuclear localisation of STAT1 whereas GBP5 expression was detected outside the nuclear membrane suggesting association with golgi in cardiomyocytes (Fig. 2e, f). Time-course studies performed in order to track the early inflammatory modulations showed that increase in *CASP1* level precedes elevation of *AIM2/NLRC4* and this occurs without alteration in *NLRP3* levels (Fig. 2g). TGFb and IL10 are recognised to exert predominantly immunosuppressive effect and regulate exaggerated immune response in human T cells and immune cells^34^. Interestingly, hiPSC-derived cardiomyocytes exhibit a swift increase in *CCL2* starting from 30 min onwards, along with increase in *TGFb* at 48h (Fig. 2h). Additionally, type I IFN stimulation by LPS and poly I:C showed differential expression of inflammatory and inflammasome marker genes. Unlike NRVCMs, hiPSC-derived cardiomyocytes showed delayed response to LPS in upregulating expression of inflammatory and inflammasome marker genes (Fig. S2c-S2h). On the other hand, despite a robust increase in inflammatory marker gene expression (*CCL2*, *CCL5*) and *CASP1*, poly I:C stimulation in hiPSC-derived cardiomyocytes failed to alter the expression of inflammasome marker genes (*AIM2*, *NLRC4*, *NLRP3*). These findings highlight species-selective differential regulation of inflammasome pathway and anti-inflammatory feedback mechanisms at play to counteract persistent inflammatory response of IFNγ in cardiomyocytes.

**Figure 2.**
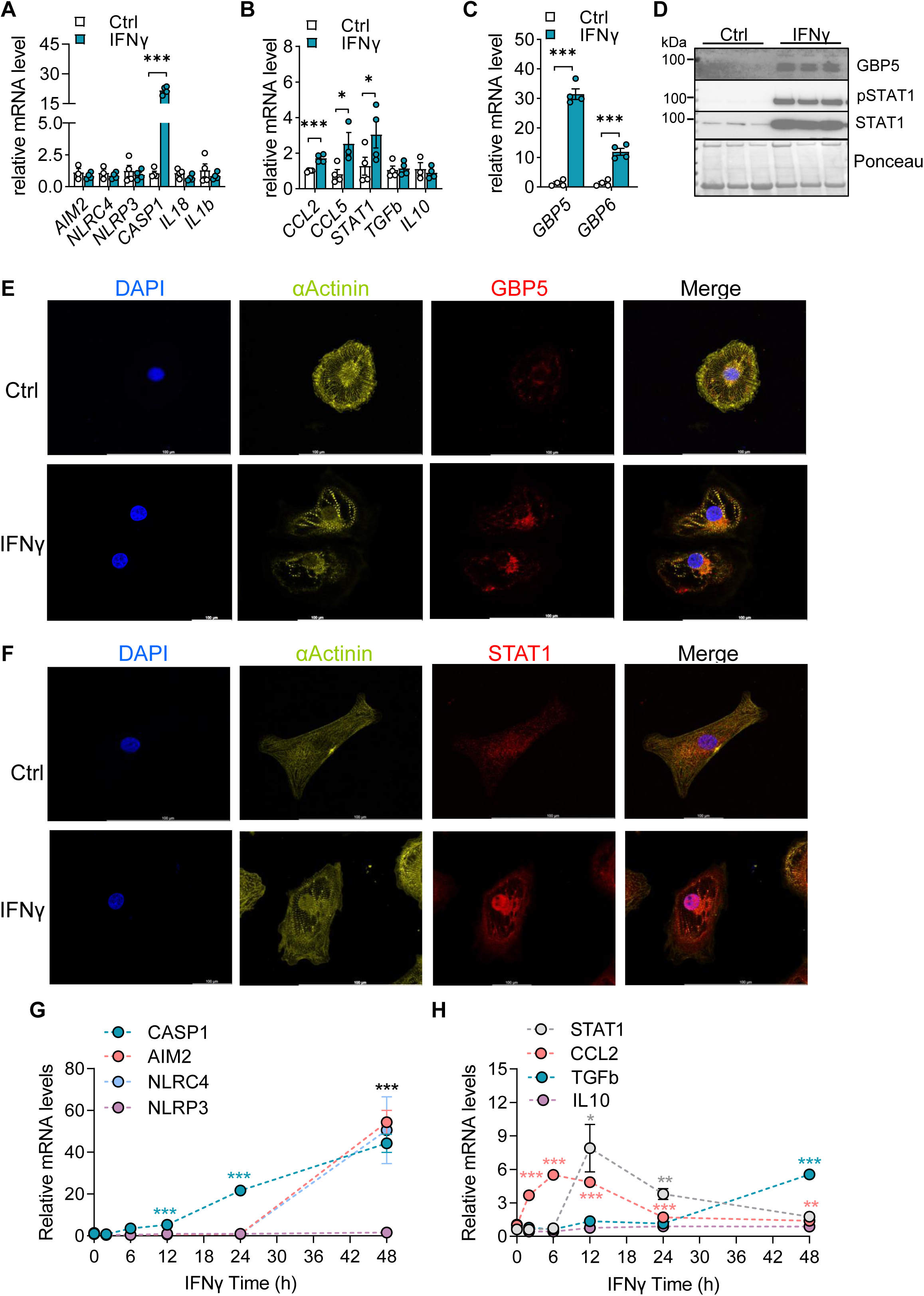
IFNγ induces GBP5/CASP1 mediated inflammasome signaling in hiPSC-derived cardiomyocytes. **A.** Expression of inflammasome markers genes upon stimulation of hiPSC-CMs with IFNγ. N=4/group. ***p<0.001, *p<0.05. **B.** Expression of inflammatory markers genes upon stimulation of hiPSC-CMs with IFNγ. N=4/group. ***p<0.001, *p<0.05. **C.** Gene expression of *GBP5* and *GBP6* upon stimulation of hiPSC-CMs with IFNγ. N=4/group. ***p<0.001. **D.** Immunoblot showing GBP5, pSTAT1 and STAT1 levels in hiPSC-CMs with IFNγ. **E.** Immunofluorescence images showing expression of GBP5 upon stimulation of hiPSC-Cardiomyocytes with IFNγ compared to controls. **F.** Immunofluorescence images showing cellular localization of STAT1 upon stimulation of hiPSC-Cardiomyocytes with IFNγ compared to controls. **G.** Time dependent effect on expression of inflammasome marker genes (CASP1, AIM2, NLRC4, NLRP3, and *CASP1*) upon stimulation of hiPSC-CMs with IFNγ. N=4/group. ***p<0.001. **H.** Time dependent effect on expression of inflammatory marker genes (STAT1, *CCL2, TGFb,* and IL10) upon stimulation of hiPSC-CMs with IFNγ. N=4/group. ***p<0.001, **p<0.01, *p<0.05.

### GBP5 deficiency attenuates IFNγ-induced CASP1 expression in cardiomyocytes

Our results in the cardiomyocytes showed that increase in *GBP5/CASP1* levels preceded elevation of *AIM2/NLRC4* in response to IFNγ stimulation. Thus, we tested the role of GBP5 deficiency in regulating IFNγ dependent inflammation and inflammasome signaling in cardiomyocytes. To address this, we subjected hiPSC-derived cardiomyocytes and NRVCMs to siRNA mediated GBP5 knock-down followed by IFNγ treatment (Fig. 3a). Downregulation of GBP5 in hiPSC-derived cardiomyocytes led to reduced *AIM2*, *NLRC4* expression, however, we found increased expression of pro-inflammatory markers (*CCL2*, *STAT1*) (Fig. 3b-d). Our data show that downregulation of GBP5 in hiPSC-derived cardiomyocytes led to increase in pro-inflammatory markers along with increased *NLRP3*, *IL18* and *IL1b* levels and a simultaneous decline in anti-inflammatory *TGFb* and *IL10* contributing to an overall pro-inflammatory regulation (Fig. 3c, 3d). On the other hand, NRVCMs with *Gbp5* knock-down showed reduced *Casp1* expression with no major effect on *Stat1* and *Ccl2* (Fig. 3e-g). However, similar to hiPSC-derived cardiomyocytes *Nlrp3* and *IL1b* expression was increased in NRVCMs (Fig. 3f). CASP7 is a downstream mediator of CASP1 signaling in the inflammasome pathway downstream signaling^35, 36^. This prompted us to determined CASP7 protein levels in the cardiomyocytes due to non-specific reactivity of available CASP1 antibodies. We found reduced CASP7 protein levels in NRVCM and trend to diminution in hiPSC-derived cardiomyocytes further suggesting a GBP5-dependent role in cardiomyocytes (Fig. 3h-k). Taken together, contrary to the previous findings from immune cells, our data highlights the paradoxical role of IFNγ wherein IFNγ-induced GBP5 exerts a cell-autonomous anti-inflammatory role in cardiomyocytes suggesting a feedback regulatory impact on inflammasome signaling.

**Figure 3.**
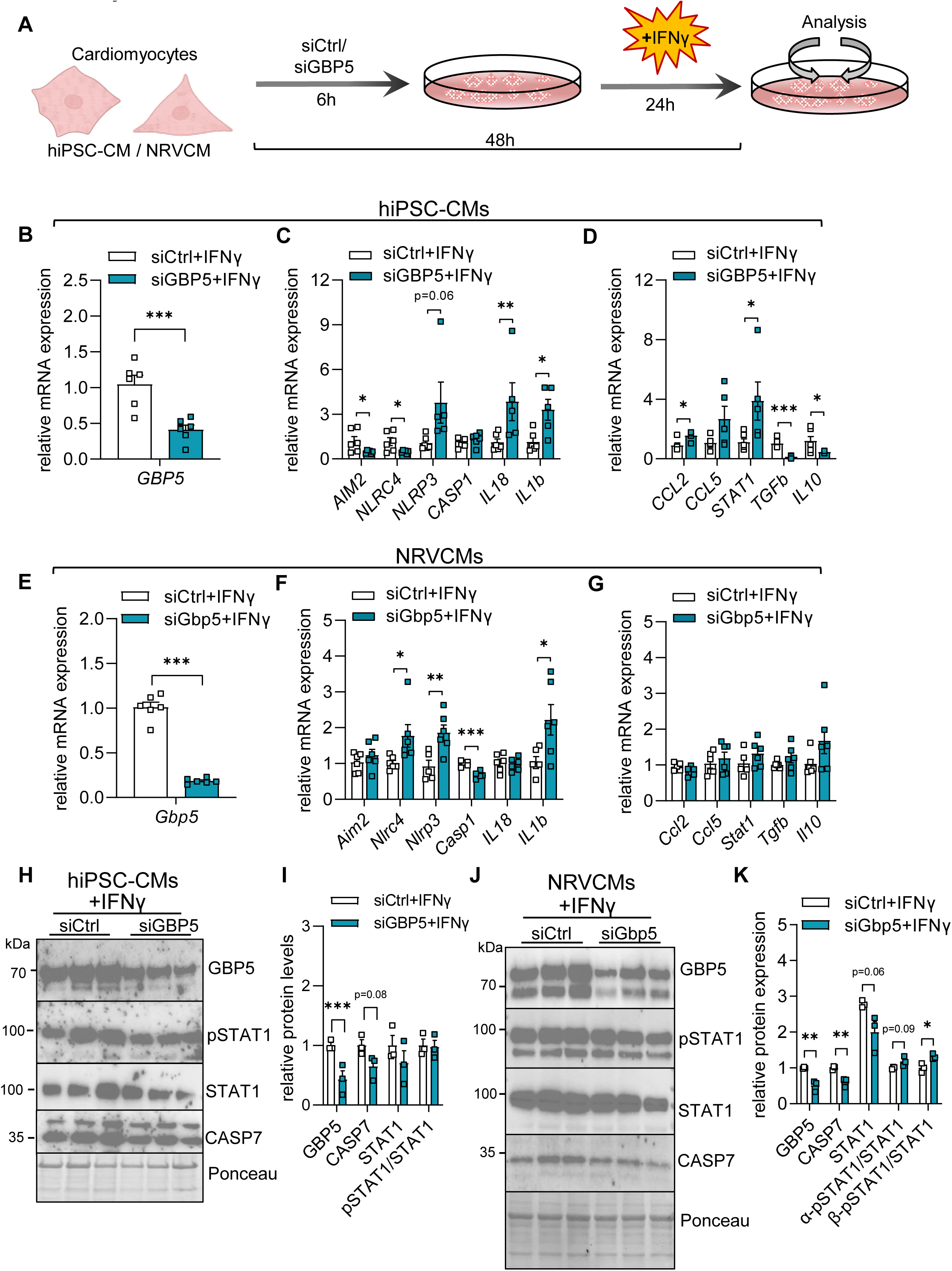
IFNγ induced CASP1 expression in cardiomyocytes is attenuated by GBP5 deficiency. **A.** Schematic representation of siRNA mediated GBP5 knock-down in hiPSC-cardiomyocytes and NRVCMs. About 75% confluent cardiomyocytes were treated with siCtrl or respective siRNA against human or GBP5 for 6h and treated with IFNγ (10ng/ml) for 24h before cell harvest. **B.** Expression of GBP5 mRNA levels after siRNA mediated knockdown of GBP5 followed by IFNγ stimulation in hiPSC-cardiomyocytes. ***p<0.001. **C.** Expression of inflammasome marker genes after siRNA mediated knockdown of GBP5 followed by IFNγ stimulation in hiPSC-cardiomyocytes. **p<0.01, *p<0.05. **D.** Expression of inflammatory marker genes after siRNA mediated knockdown of GBP5 followed by IFNγ stimulation in hiPSC-cardiomyocytes. **E.** Expression of GBP5 mRNA levels after siRNA mediated knockdown of GBP5 followed by IFNγ stimulation in NRVCMs. ***p<0.001. **F.** Expression of inflammasome marker genes after siRNA mediated knockdown of GBP5 followed by IFNγ stimulation in NRVCMs. **p<0.01, *p<0.05. **G.** Expression of inflammatory marker genes after siRNA mediated knockdown of GBP5 followed by IFNγ stimulation in NRVCMs. **H.** Immunoblot showing protein levels of GBP5, CASP7, pSTAT1 and STAT1 after siRNA mediated knockdown of GBP5 followed by IFNγ stimulation in hiPSC-cardiomyocytes. **I.** Quantification of the immunoblots shown in Fig 3H. ***p<0.001. **J.** Immunoblot showing protein levels of GBP5, CASP7, pSTAT1 and STAT1 after siRNA mediated knockdown of GBP5 followed by IFNγ stimulation in hiPSC-cardiomyocytes. **K.** Quantification of the immunoblots shown in Fig 3J. *p<0.05.

### GBP5 expression attenuates the IFNγ-induced inflammation and inflammasome signaling in cardiomyocytes

In order to investigate the direct effect of GBP5 expression in regulating cardiomyocytes inflammation we generated human GBP5 expression construct. Immunofluorescence analysis in hiPSC-derived cardiomyocytes expressing human GBP5 showed GBP5 localisation around nucleus suggesting golgi association (Fig. 4a) similar to endogenous GBP5 expression observed with IFNγ stimulus (Fig. 2e). We standardized the GBP5 adenovirus transduction conditions to attain comparable expression of GBP5 between hiPSC-derived cardiomyocytes expressing GBP5 (5 MOI) and IFNγ stimulation (Fig. 4b, 4c). Importantly, IFNγ stimulation failed to further increase GBP5 expression in hiPSC-derived cardiomyocytes transduced with GBP5 suggesting an anti-inflammatory role of GBP5 (Fig. 4c, 4e). Further analysis showed that GBP5 overexpression in cardiomyocytes showed no effect on *AIM2*, *NLRC4* levels whereas expression of inflammasome markers *CASP1 and IL18* was reduced in the absence or presence of IFNγ stimulation, respectively (Fig. 4d). However, GBP5 expression attenuated the increase in IFNγ-induced inflammasome markers (*AIM2*, *NLRC4*, *CASP1*, *IL1b*, *IL18*). Similarly, despite an increase in basal *TGFb* expression by increased GBP5 levels, IFNγ-induced *TGFb* levels failed to escalate suggesting a feedback mechanism in hiPSC-derived cardiomyocytes (Fig. 4f). Furthermore, GBP5 expression attenuated cellular inflammation with decreased mRNA levels of *CCL2* and *STAT1* (Fig. 4e, 4f).

**Figure 4.**
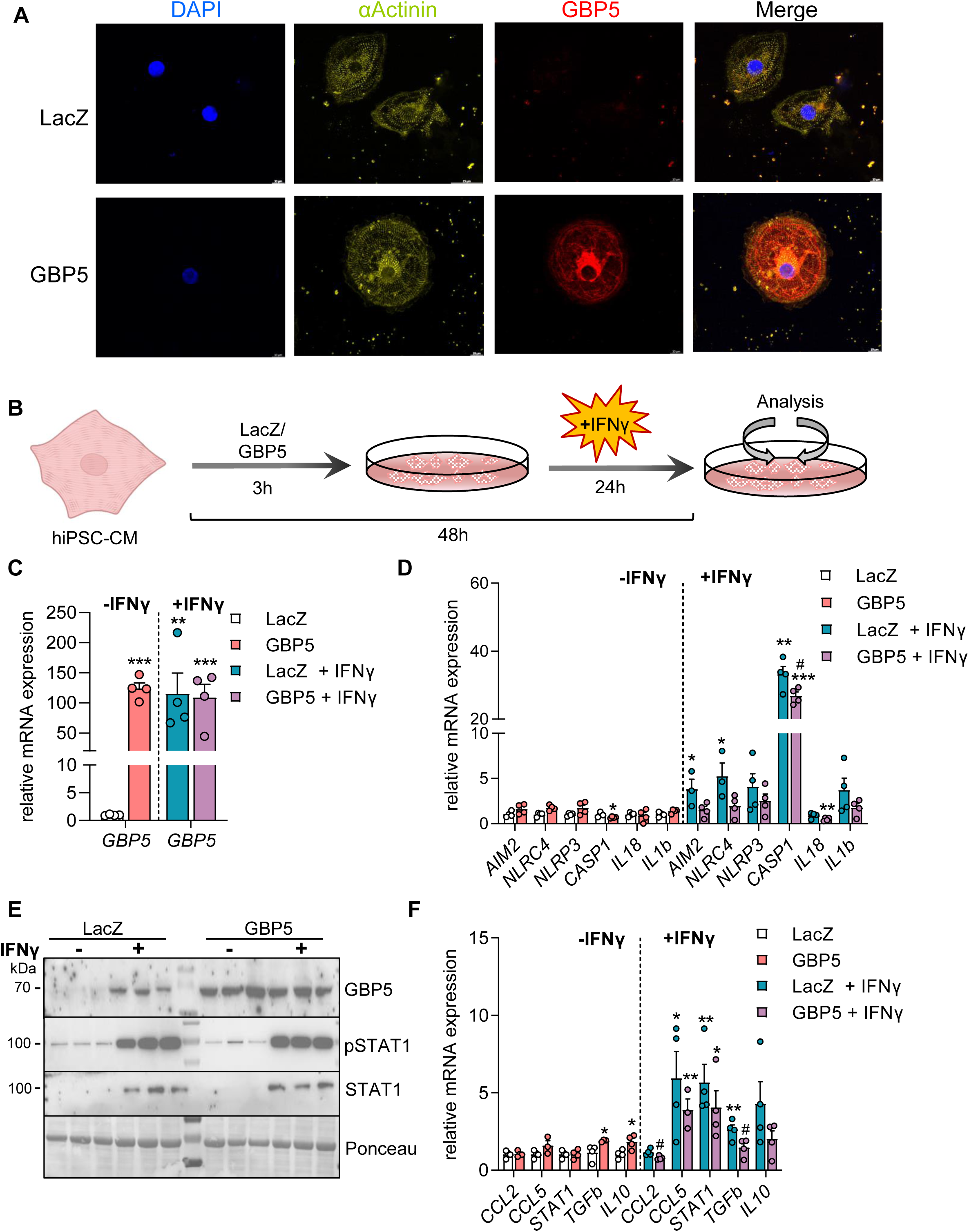
GBP5 expression attenuates the IFNγ-induced inflammatory and inflammasome signaling in hiPSC-derived cardiomyocytes. **A.** Immunofluorescence images showing GBP5 expression in hiPSC-CMs upon stimulation with IFNγ (10ng/ml) compared to LacZ controls. **B.** Schematic representation of GBP5 expression in hiPSC-cardiomyocytes. About 75% confluent cardiomyocytes were transduced with adenovirus expressing GBP5 or LacZ control and treated with IFNγ (10ng/ml) before cell harvest. **C.** Effect on GBP5 mRNA levels after adenovirus mediated expression of GBP5 in hiPSC-CMs in the absence and presence of IFNγ. N=4/group. * compared to LacZ control (- IFN γ), ^#^ compared to LacZ control (+IFN γ). ***p<0.001, *p<0.05, ^###^p<0.001. **D.** Effect on inflammasome marker genes after adenovirus mediated expression of GBP5 in hiPSC-CMs in the absence and presence of IFNγ. N=4/group. * compared to LacZ control (-IFN γ), ^#^ compared to LacZ control (+IFNγ). **p<0.01, *p<0.05, ^#^p<0.05. **E.** Immunoblot showing expression of GBP5, pSTAT1, STAT1 protein levels in hiPSC-CMs upon adenovirus mediated expression of GBP5 in the absence and presence of IFNγ. N=3/group. **F.** Effect on inflammatory marker genes after adenovirus mediated expression of GBP5 in hiPSC-CMs in the absence and presence of IFNγ. N=4/group. * compared to LacZ control (-IFN γ). **p<0.01, *p<0.05, ^#^p<0.05.

Next, we determined the effect of human GBP5 expression in NRVCMs (Fig. 5a). Interestingly, human *GBP5* overexpression in NRVCM attenuated the endogenous as well as IFNγ-stimulated *Gbp5* expression in NRVCMs reiterating the potential anti-inflammatory role of GBP5 in cardiomyocytes (Fig. 5b, 5c). Corroborating the findings from hiPSC-derived cardiomyocytes, human *GBP5* expression in NRVCMs attenuated IFNγ-induced expression of inflammatory marker genes (*Ccl2* and *Stat1*) (Fig. 5d, 5e) and led to reduced expression inflammasome signaling (*Aim2*, *Nlrc4*, *Nlrp3*) (Fig. 5f). Additionally, similar to hiPSC-derived cardiomyocytes IFNγ-induced *Casp1* and *Tgfb* expression was not further altered with increase in *GBP5* levels suggesting a feedback regulation in NRVCMs. These findings show that GBP5 expression in cardiomyocytes attenuates the IFNγ-driven pro-inflammatory signaling cascade. The anti-inflammatory effects of GBP5 expression in cardiomyocytes suggest the potential of GBP5 to act as an inflammatory brake on IFNγ stimulated inflammasome pathway.

**Figure 5.**
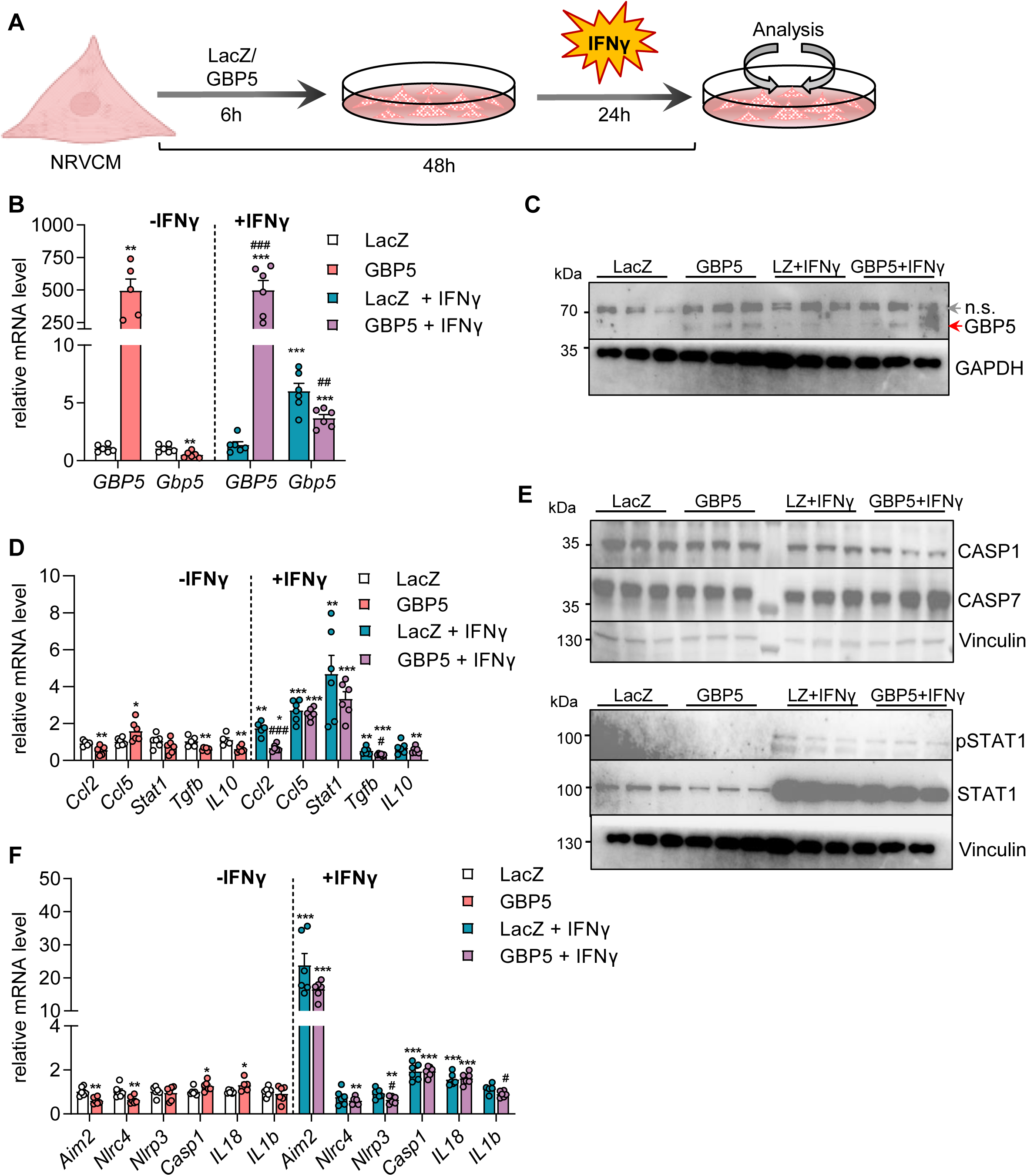
GBP5 attenuates the IFNγ-induced inflammatory and inflammasome signaling in NRVCMs. **A.** Schematic representation of GBP5 expression in NRVCMs. About 75% confluent cardiomyocytes were transduced with adenovirus expressing GBP5 or LacZ control and treated with IFNγ (10ng/ml) before cell harvest. **B.** Effect on GBP5 mRNA levels after adenovirus mediated expression of human *GBP5* and rat *Gbp5* in NRVCMs in the absence and presence of IFNγ. N=6/group. * compared to LacZ control without IFNγ, ^#^ compared to LacZ control with IFNγ. ***p<0.001, **p<0.01, ^##^p<0.01, ^###^p<0.001. **C.** Immunoblot showing adenovirus mediated expression of GBP5 in NRVCM in the absence and presence of IFNγ. GBP5 and the non-specific (n.s.) band are marked. **D.** Effect on inflammatory marker genes after adenovirus mediated expression of GBP5 in NRVCMs in the absence and presence of IFNγ. N=4/group. * compared to LacZ control without IFNγ, ^#^ compared to LacZ control with IFNγ. **p<0.01, *p<0.05, ^#^p<0.05, ^###^p<0.001. **E.** Immunoblot showing expression of CASP1, CASP7, pSTAT1, and STAT1 protein levels in NRVCMs upon adenovirus mediated expression of GBP5 in the absence and presence of IFNγ. N=3/group. **F.** Effect on inflammasome marker genes after adenovirus mediated expression of GBP5 in NRVCMs in the absence and presence of IFNγ. N=4/group. * compared to LacZ control without IFNγ, ^#^ compared to LacZ control with IFNγ. **p<0.01, *p<0.05, ^#^p<0.05.

### GBP5 deficiency augments IFNγ-induced cell death in cardiomyocytes

Inflammatory signaling elicited due to IFNγ often modulates cell proliferation and apoptosis^37^. In cardiomyocytes, the detrimental effects of heightened inflammatory state can be captured by measuring the cardiomyocyte contractile function. Our data show that cardiomyocytes exhibited reduced amplitude of contraction, reduced contraction- and relaxation-velocity along with increased beats per minute suggesting overall impaired cardiomyocyte contractility in the presence of IFNγ stimulation (Fig. 6a-d, S3a-S3c). Impaired cardiomyocyte contractility can alter mitochondrial respiration. In this regard we found that IFNγ stimulation significantly reduced basal oxygen consumption rate (OCR) in cardiomyocytes (Fig. 6e). Additionally, we determined the mitochondrial oxidative function by quantifying mitochondrial OXPHOS proteins, namely NDUFB8 (Complex I), SDHB (Complex II), CoI (Complex IV), UQCRC2 (Complex III), and ATP5A (Complex V). IFNγ stimulation in cardiomyocytes results in reduced levels of Complex I, Complex II, Complex III and Complex IV suggesting a compromised oxidative potential (Fig. 6f, 6g). To note, the cellular metabolic activity as determined by MTT assay was not altered in the IFNγ stimulated cardiomyocytes (Fig. S4a). However, downregulation of GBP5 by siRNA mediated knockdown augmented cell death in cardiomyocytes stimulated with IFNγ indicating culmination of the deleterious effect of exaggerated inflammation onto cardiomyocyte apoptosis (Fig. 6h-k). To the contrary, GBP5 overexpression in NRVCMs did not alter cardiomyocyte cell death (Fig. 6l, 6m).

**Figure 6.**
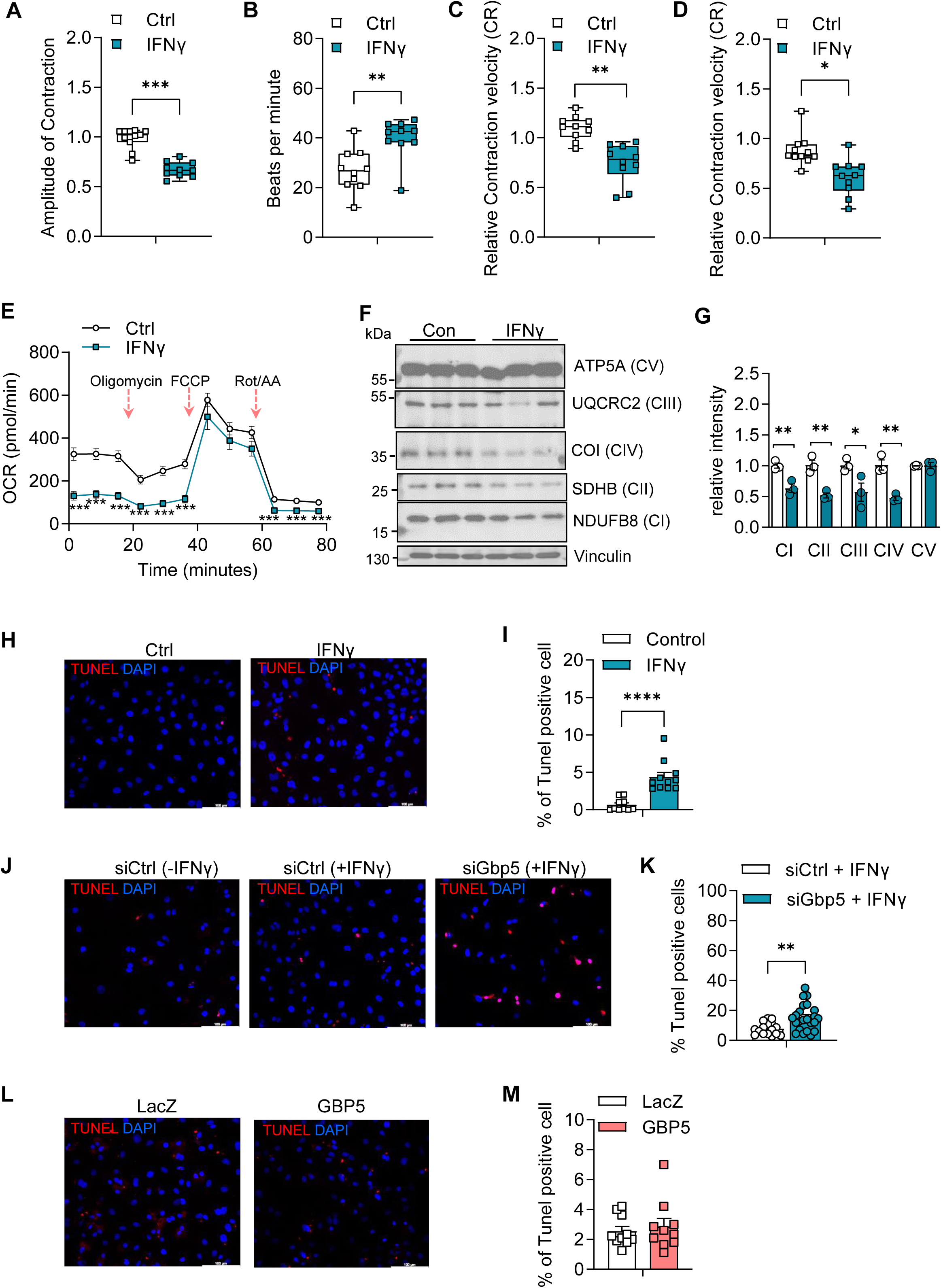
GBP5 deficiency augments IFNγ-induced cell death in cardiomyocytes. **A-D**. Effect of IFNγ on cardiomyocyte contractile function assessed by cardiomyocyte contractility assay in NRVCMs upon stimulation with IFNγ (10ng/ml) compared to controls. N=12 per group **E**. Mitochondrial respiration was determined by Cell Mito Stress Assay in cardiomyocytes upon IFNγ (10ng/ml) using Seahorse XF analyzer. ***p<0.0001, *p<0.05. **F**. Effect on mitochondrial oxidative phosphorylation was determined by immunoblotting NDUFB8 (Complex I), SDHB (Complex II), CoI (Complex IV), UQCRC2 (Complex III), ATP5A (Complex V) after IFNγ (10ng/ml) stimulation in cardiomyocytes. **G**. Quantification of the immunoblot shown in Fig. 6F. **p<0.01, *p<0.05. **H-I**. Effect of IFNγ (10ng/ml) stimulation on cardiomyocyte cell death assessed by quantifying DNA damage using TUNEL assay. ***p<0.0001. **J-K**. Effect of siRNA mediated GBP5 knockdown followed by IFNγ (10ng/ml) stimulation on cardiomyocyte cell death assessed by quantifying DNA damage using TUNEL assay. **p<0.01. **L-M**. Effect of GBP5 overexpression on cardiomyocyte cell death assessed by TUNEL assay based DNA damage quantification.

**Figure 7.**
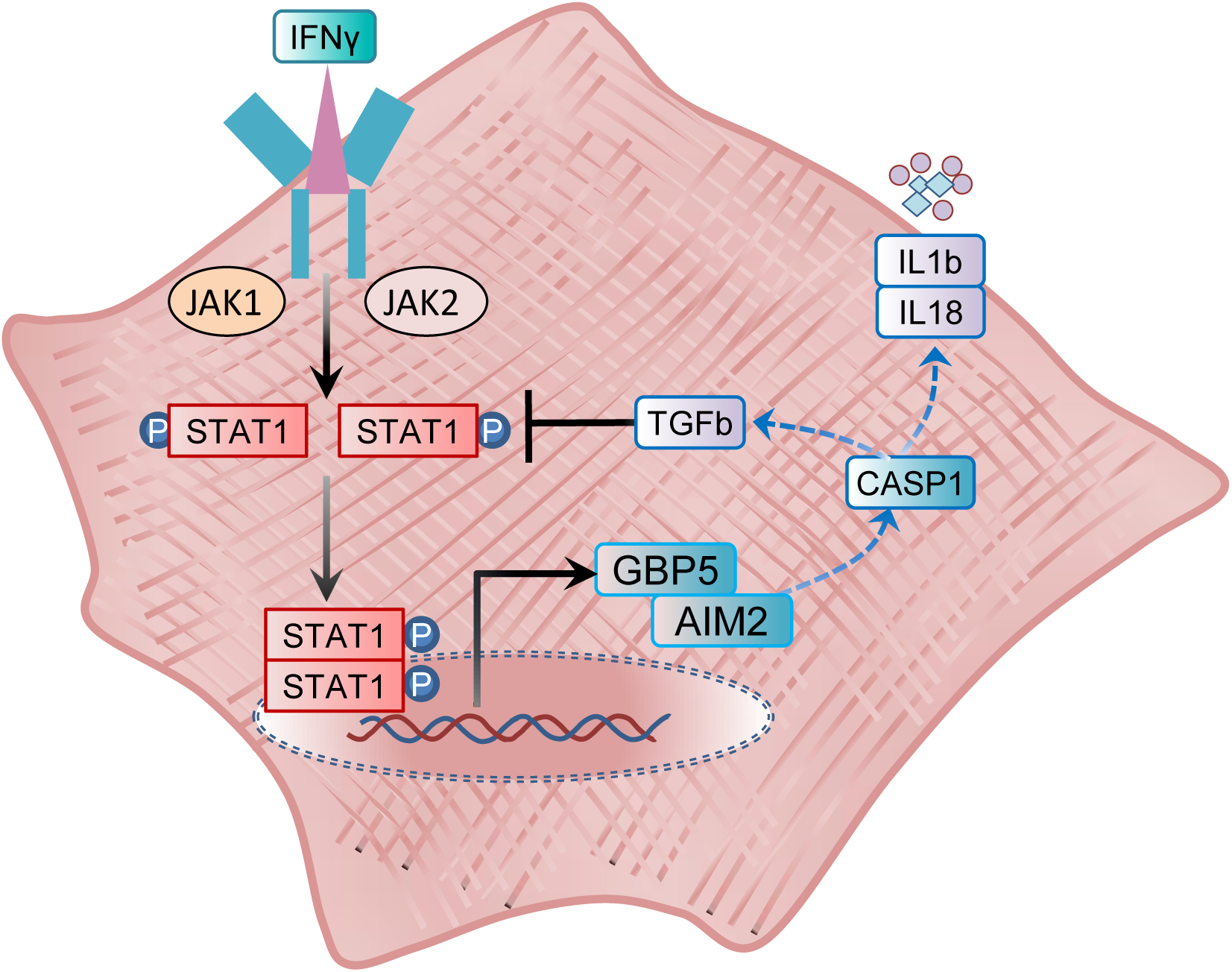
Schematic representation of the interplay between pro- and anti-inflammatory signaling of IFNγ and GBP5 in hiPSC-derived cardiomyocytes. The schematic figure shows that IFNγ initiates pro-inflammatory effect in hiPSC-derived cardiomyocytes by inducing STAT1, GBP5, AIM2 and CASP1 expression. This in turn follows a feedback inhibitory effect on IFNγ signaling via GBP5/TGFb on STAT1 levels creating cellular rheostat by attenuating the excessive pro-inflammatory response of IFNγ.

## Discussion

Diverse effects of IFNγ signaling have been reported in the myocardium of patients with ischemic, non-ischemic and anti-cancer drugs associated cardiomyopathies. Our understanding of these underlying inflammatory components has emerged primarily from studies elucidating function of immune cells within myocardium whereas the role of cardiomyocytes remains unclear. Here we identify cardiomyocytes as immunocompetent cell displaying robust upregulation of inflammasome signaling via *GBP5/CASP1* mediated pathway followed by *AIM2/NLRC4* in response to IFNγ. We show that by virtue of *GBP5/TGFb* axis, cardiomyoctes orchestrate a parallel anti-inflammatory feedback regulation to combat IFNγ-induced inflammatory signaling.

Multiple independent studies highlight the role of IFN induced alteration in NLRP3 and AIM2 inflammasomes in the pathophysiology of cardiovascular disease. While innate immune response is a hallmark of sterile inflammation in various cardiomyopathies, inflammasome acts as an intracellular sensor responding to DAMPs upon tissue damage. Evidence from multiple transgenic mouse models and drugs targeting IL1b, NLRP3 has established their role in inflammasome biology studying the pathophysiology of cardiac dysfunction^38^. Increased expression of NLRP3 is associated with maladaptive cardiac remodelling and fibrosis in myocardial infarction and atherosclerosis. On the other hand, elevated AIM2 expression has gained interest in the past 8 years, especially in the context of cardiovascular disease as a key driver of atherosclerosis and DNA damage^39, 40^. It is noteworthy that AIM2 expression has also been shown to be upregulated in the failing heart of patients with ischemic and dilated cardiomyopathy with unaltered NLRP3 levels suggesting a key driver of human cardiovascular disease^15^. NLRP3 and AIM2 execute context specific role because NLRP3 is activated by DAMPs such as ROS, mtDNA whereas AIM2 is specifically activated by cytoplasmic dsDNA. In this regard, results from our comparative study show that cardiomyocytes upregulate *NLRP3* in response to type I IFN stimuli whereas type II IFN activation resulted in selective upregulation of *AIM2*. These findings indicate cell-type specific differential regulation of AIM2 inflammasome in cardiomyocytes and align with previous research suggesting increased ROS and mtDNA damage by LPS and PolyI:C stimulation involved in modulating NLRP3 levels^41, 42^. Concordant to the increase in AIM2 levels we identified increased DNA damage and impaired mitochondrial respiration and oxidative function in cardiomyocytes stimulated with IFNγ.

The inflammasomes activates CASP1 to release pro-inflammatory cytokines such as IL1b and IL18. The Canakinumab Antiinflammatory Thrombosis Outcome Study (CANTOS), a randomized, double-blind trial on 10,061 recruited patients demonstrates the proof of concept that inflammasome targeting via IL1b innate immunity pathway plays a significant role in lowering major cardiovascular events^43^. Despite similarities, clinical studies and transgenic models highlight species-specific and cell-specific divergence in non-canonical Caspases, inflammasome complexity, differential transcriptional control of inflammasome signaling^44, 45^. Our comparative analysis from NRVCMs and hiPSC-derived cardiomyocytes also highlight the species differences in the inflammation and inflammasome executors and regulators within cardiomyocyte in response to IFNγ that warrants further validation before direct interpretation of findings from rodent models to human cardiac inflammasome regulation.

Cardiomyocytes respond to danger signals by cytokines and chemokines secretion, and subsequent recruitment of leucocytes and cell-surface adhesion molecules^46^. Among these, TGFβ and IL10 are potent anti-inflammatory cytokines that are known to alter the dynamics of IFNγ secretion from T cells and NK cells^27^. TGFβ is a multifunctional polypeptide cytokine and similar to IFNγ exerts dual and paradoxical effects on fibrosis and immune regulation at cellular and intercellular level upon cardiac injury^47^. Early TGFβ inhibition in myocardial infarction increases mortality, whereas late TGFβ inhibition attenuates adverse cardiac remodeling. Lethal consequences of TGFβ signaling deletion in mice caused by enormous and uncontrolled inflammation underscores a critical role in inflammatory circuits^48^. These findings are in congruence to the studies identifying anti-inflammatory properties of traditionally presumed pro-inflammatory IFNγ regulating innate immune response early after myocardial infarction and myocardial hypertrophy^7, 8^. TGFβ can suppress IFNγ-STAT1 activation via regulating canonical and non-canonical pathway. In cardiomyocytes, TGFβ/SMAD signaling in cardiomyocytes can modulate miR486 levels to inhibit STAT1 expression or activate non-canonical TGFβ-activated kinase 1(TAK1) pathway to modulate STAT1 levels^49, 50^.

Our study with GBP5 knockdown and overexpression identify an intercellular feedback regulation of IFNγ driven inflammasome pathway via downregulation of *STAT1* expression driven by *GBP5/TGFb* expression. We show that GBP5 knockdown in hiPSC-derived cardiomyocytes upregulates STAT1 expression, whereas GBP5 overexpression failed to increase STAT1 levels. This suggests that pro-inflammatory IFNγ signaling in cardiomyocytes exists in coherence with a parallel anti-inflammatory pathway exerting a feedback regulation via TGFβ. In addition, these observations warrant further detailed analysis of intercellular cross-talk studies in vivo to deconvolute the contribution of cardiomyocyte in IFNγ stimulated inflammasome signaling.

Of note, GBP5 has been shown to exert disease-selective regulatory role in inflammation. Activation of autophagy by GBP5 downregulation was shown to be beneficial against LPS-induced lung injury^51^. Haque et. al. showed that GBP5 exhibits anti-inflammatory role in the context of rheumatoid arthritis^52^. GBP5 has been largely studied in regulating innate immune response against bacteria, viruses and parasite by NLRP3 driven canonical as well as non-canonical inflammasome signaling^53–55^. On the other hand, GBP5 suppression by nuclear receptor ERRγ has been shown to be protective against LAD-induced myocardial infarction^20^. Importantly, GBP5 is a marker of immuno-hot tumors and upregulated in the myocardium of a subset of cancer patients diagnosed with immune check point inhibitors-induced myocarditis^18, 56^. High infiltration of CD8+T cells in the myocardium of few patients treated with immune check point inhibitors thus renders cardiomyocytes vulnerable to a microenvironment of increased inflammation and IFNγ signaling^57^. Overall, broader consensus recognizes a pro-inflammatory role of GBP5 enhancing NLRP3 inflammasome. However, our findings show that GBP5 levels at the basal state are extremely low in cardiomyocytes but are highly inducible upon IFNγ stimulation. Additionally, upregulation of GBP5 followed by AIM2 and CASP1 provides insights into an immunocompetent role of cardiomyocytes. Our GBP5 knockdown and overexpression studies identify an anti-inflammatory role of GBP5 within cardiomyocytes. Nevertheless, the intracellular and intercellular communication in the myocardium due to altered GBP5 levels in the cardiomyocytes remains undefined and is plausible that the disease severity and causal factors in turn alter the inflammation regulatory trajectories in vivo. Identifying alteration in GBP5 levels in transgenic mouse models and drugs targeting IL1b, NLRP3 and AIM2 will further advance our understanding of inflammasome biology in the context of pathophysiology of cardiac dysfunction^38, 58^.

Taken together, driving the paradoxical actions of IFNγ as an effector of inflammasome response and regulator of cytokine signaling, the duration and the context of cardiac damage play a decisive role in the pathogenesis of heart failure. Thus, GBP5/AIM2/CASP1/TGFb cascade in the cardiomyocytes serves as a physiological adaptive pro- and anti-inflammatory pathway executed in parallel that may enable rheostat generation in the myocardium towards IFNγ response against sterile inflammation.

## Material and Methods

### Isolation of neonatal rat ventricular cardiomyocytes (NRVCMs)

NRVCMs were isolated using the MACS® Cell Separation System (Miltenyi Biotec B.V. & Co. KG, Bergisch Gladbach, Germany) together with the Neonatal Cardiomyocyte Isolation Kit (Miltenyi Biotec B.V. & Co. KG, Bergisch Gladbach,Germany). In brief, left ventricles from 1-3 days old Wistar rats (JANVIER LABS, Lyon, France) were harvested and washed 2 to 3 times in ADS buffer containing 116.3 mM NaCl, 19.7 mM HEPES, 9.4 mM NaH_2_PO_4_, 5.5 mM Glucose, 5.3 mM KCl and 0.8 mM MgSO_4_, pH 7.4. The hearts were cut into pieces to a homogeneous mixture, with the ADS being aspirated beforehand. Shredded hearts were distributed equally to C-Tubes, where 5 mL of Enzyme Mix I+II (containing enzyme P, A and D together with buffer X and Y) was added on top. C-Tubes were put on device for digestion, centrifuged and strained through a cell strainer. For cardiomyocyte isolation, the supernatant was discarded and the pellet was resuspended with 60 µL PEB buffer, containing 0.5% BSA and 2 mM EDTA. On top 20 µL of isolation mix and Anti-red blood Cell Microbeads were added to facilitate the isolation process. After cardiomyocytes were isolated through the MACS Magnet, cells were re-suspended in culture media (DMEM:F12 media containing 10% FCS, 1% penicillin, 1% streptomycin and 1% L-glutamine, Gibco Thermo Fisher Scientific, Waltham, Massachusetts, U.S.), counted and seeded into culture plates.

### Culture of human iPSC-derived cardiomyocytes

Human iPSC-derived cardiomyocytes (hiPSC-CMs) were generated by the Stem Cell Core Facility (Universitätsklinikum Heidelberg). Briefly, hiPSCs were thawed onto Matrigel-coated 6-well plates in hE8 medium supplemented with ROCK inhibitor (1:1000), centrifuged at 300 × g for 3 min, resuspended in hE8/ROCK inhibitor, and cultured at 37 °C and 5% CO_2_. Differentiation into cardiomyocytes was induced using a small-molecule-based protocol, followed by a glucose starvation phase from day 10 to day 12 to enrich for metabolically resilient cardiomyocytes. hiPSC-CMs were maintained until day 40–50 in RPMI 1640 containing L-ascorbic acid 2-phosphate (2.11 g/L) and human serum albumin (5 g/L), supplemented with B27 (Gibco), with medium changes every 24 h before use in experiments.

### Cloning of human GBP5

Human GBP5 was cloned using cDNA from Human Umbilical Vein Endothelial Cells (HUVECs) stimulated with IFNγ (10ng/ml). Target sequences were amplified via specific Fishing PCR using primers: 5’-GGGGACAAGTTTGTACAAAAAAGCTGGCACCATGGCTTTAGAGATCCAC-3’ and 5’-GGGGACCACTTTGTACAAGAAAGCTGGGTCGCCTTAGAGTAAAACACATGG-3’ in pDonR221 gateway cloning vector (Thermo Scientific) following the manufacturer’s instructions. Cycling conditions were: initial denaturation (98°C, 30 s), followed by 39 cycles of denaturation (98°C, 10 s), annealing (55°C, 30 s), extension (72°C, 1 min), and a final extension (72°C, 10 min). PCR products were purified with the NucleoSpin Plasmid Kit (Macherey-Nagel) as per manufacturer instructions and cloned into pDonR221 entry vector using Gateway BP Clonase II. Positive clones were isolated via NucleoSpin Plasmid MiniPrep Kit. Entry clones underwent LR recombination into pAd/CMV/V5-DEST destination vector and verified by Sanger sequencing (Azenta GeneWiz).

### Generation of recombinant adenovirus and titre

Validated plasmids were digested with PacI restriction enzyme and transfected into HEK293A cells using Lipofectamine 2000 to produce respective protein-expressing adenovirus. Adenoviral titration was performed in HEK293A cells by staining with fluorescent anti-Hexon antibody and IFU/mL was determined. A β-galactosidease-V5-encoding adenovirus (Ad-LacZ) served as a control.

### Cell culture and treatment

Isolated NRVCM were cultured in cell culture plates coated with collagen (50µg/ml). Cell culture plates coated with collagen were incubated for 1h at 37°C at 5% CO2, aspirated and washed using water for injection (WFI, Gibco Thermo Fisher Scientific, Waltham, Massachusetts, U.S.). NRVCMs were seeded and allowed to adhere for 24 h. After 24 h, cells were briefly washed with DBPS. Overexpression and knockdown studies for gene expression and immunoblotting was achieved using adenovirus-mediated transduction for 48h as mentioned above. NRVCMs were treated with IFNγ (10ng/ml); Poly I:C (1µg/mL) and LPS (100ng/ml) for indicated time points mentioned in the respective figure legends. After 48h, cells were washed with PBS and harvested for RNA and protein isolation as described below.

### GBP5 knockdown by siRNA silencing

To obtain GBP5 knockdown, siRNAs specific to human or rat guanylate-binding protein 5 (GBP5) were purchased from ThermoFisher Scientific (Cat. Number, rat: 4390771; Cat. Number, human: 4392420). Briefly, 60 – 80 % confluent cells were used for transfection. On the day of transfection, siRNAs were diluted in DMEM:F12 to the desired final concentration of 100 pmol. Separately, Lipofectamine™ RNAiMAX was also diluted in DMEM:F12 and incubated for 5 min at RT. The diluted siRNA and RNAiMAX solutions was combined and incubated for 15 min at RT to allow formation of siRNA-lipid complexes. NRVCMs were incubated with the transfection mixture at 37 °C in a humidified CO_2_ incubator for 6 h. After this incubation period, the medium was carefully replaced with fresh complete growth medium and cells were allowed to grow for an additional 18 h under standard culture conditions. NRVCMs were stimulated with IFNγ (10 ng/mL) and harvested for downstream experimental analyses.

### MTT Assay to assess cell viability

NRVCMs were cultured in 24 well plates and treated with different concentration of IFNγ to assess the effect on cell viability. Treated cells were incubated for 48h before the MTT assay. After a media change and PBS wash, fresh media, containing 10% MTT labelling reagent (Roche Applied Sciences, Penzberg, Germany), was added to the wells. After 4h of incubation at 37°C, 250µL MTT solubilization solution was added on top of the volume in each well. To allow complete formation of formazan crystal, cells were kept at 37°C overnight. The amount of violet formazan crystals formed was determined by spectrophotometric absorbance using a Tecan Spark microplate reader (Männedorf, Switzerland) at 550 nm - 600 nm, along with reference wavelength of >650 nm. The percentage of viable cells was calculated in relation to the respective controls.

### RNA isolation and quantitative real-time PCR

Total RNA was isolated using TRIzol reagent (Invitrogen, Carlsbad, U.S.), according to the manufacturer’s protocol. After RNA isolation, a total of 1µg RNA was used for cDNA synthesis using the iScript™ cDNA Synthesis Kit (Bio-Rad, Feldkirchen, Germany). For the qRT-PCR a 1:10 diluted cDNA was used together with SYBR Green real-time PCR master mix (Thermo Fisher Scientific, Waltham, Massachusetts, U.S.) on a LightCycler 480 real-time PCR system from Roche (Penzberg, Germany). Expression of RPL32 was used as an internal standard for quantification. All experiments in NRVCMs and hiPSC-CMs were performed with n = 6 (hexaplicate) or n = 4 (quadruplicate) biological replicates per experiment and repeated independently at least two-three times.

### Cardiomyocyte Contractility Assay

To evaluate alterations in contractile function, cardiomyocytes were subjected to a microscopy-based contractility assay. To assess contractility, cardiomyocytes (NRVCMs) were seeded in black 96 well plates with a glass bottom (ThermoFisher Scientific/USA). After adherence, cells were treated according to the experimental set up. 24 h before end of the treatment, cells were treated with 200 nM Tetramethylrhodamine, methyl ester (TMRM) (ThermoFisher Scientific/USA), incubated for 30 min at 37 °C with 5 % CO_2_. Thereafter, cells were washed three times with cell culture media and incubated for 24 h to enable the intercalation of the dye into the cells. Contractile function was measured using a ZEISS Axio Observer microscope, acquiring 1500 images at a frame rate of 50 frames per second. TMRM fluorescence was excited at 548 nm and emission was collected at around 574 nm. For analysis the CCA .tif image stacks were first resized using a custom Python script to ensure compatibility with the CCA v1.0 MATLAB analysis software. For data organization, a Job File was generated from the microscopy data. This Job File is a plain-text document that lists the directories containing the image stacks to be analysed, which is particularly advantageous for high-throughput experiments involving large numbers of folders. To streamline this process, CCA v1.0 provides an automated feature that recursively scans for subfolders containing .tif images, displays them within the graphical user interface (GUI), and compiles the results into a Job File. Subsequently, a configuration file was selected to define the analysis parameters. In this study, the “Frame-Frame-subtraction” configuration was employed, wherein CCA calculates the difference in pixel intensity between consecutive frames. Changes in pixel intensity are interpreted as indicators of cell movement between frames. Once the Job File and configuration file were specified, CCA v1.0 proceeded with the automated analysis of the image stacks. The software computes contractility signals, including phase-averaged contraction peaks annotated with extracted parameters, as well as velocity signals. The resulting data can be exported as .csv files for subsequent statistical analysis. The CCA provides four quantitative parameters for evaluating cardiomyocyte contractile properties: T_Peak (time to peak contraction), D_Peak (duration of peak contraction), contraction velocity (rate of cell shortening), and relaxation velocity (rate of cell lengthening). These metrics enable detailed characterization of the kinetics and dynamics of cardiomyocyte contraction and relaxation.

### TUNEL Assay

Apoptosis due to cell death was assessed using the TUNEL in situ apoptosis detection kit (Elabscience, Houston, Texas, USA), which labels DNA strand breaks by enzymatic incorporation of fluorescently labeled nucleotides at free 3’-OH termini via terminal deoxynucleotidyl transferase (TdT). NRVCM were seeded on µ-slides and subjected to adenoviral transduction for overexpression or transfected for GBP5 silencing. After treatment, cells were fixed with 4% paraformaldehyde (PFA) in PBS for 20 min at RT, followed by a single PBS wash. Fixation was then repeated with 4% PFA, after which cells underwent three washes with PBS (5 min each). Cells were permeabilized with 0.2% Triton X-100 in PBS at 37°C for 10 min, followed by three additional PBS washes (5 min each).

Each experiment included negative and positive control wells prepared as follows: Positive controls were incubated with 100 µL of 1x DNase I buffer for 5 min at room temperature followed by treatment with 100 µL DNase I working solution (200 U/mL) for 10 to 30 min at 37°C. Slides were subsequently washed three times with PBS (5 min each). Negative controls received 100 µL of 1x DNase I buffer for 5 min at room temperature and an additional 10 to 30 min incubation in the same buffer at 37°C, followed by identical PBS washes. Labelling working solution were prepared according to the number of cells to be stained, the respective volumes are shown below.

### Preparation of Labeling Working Solution

| Component | Positive control/ Experimental group | Negative control |
| --- | --- | --- |
| TdT Equilibration Buffer | 35μL | 40μL |
| Labeling Solution | 10μL | 10μL |
| TdT Enzyme | 5μL | 0μL |

As the first step, 100 µL of TdT Equilibration Buffer was added to each sample and incubated at 37°C for 10-30 min. Following this, 50 µL of the Labeling Working Solution was added to each well and incubated for 60 min in the dark, within a humidified chamber. The µ-slides were then washed three times with PBS, each wash lasting 5 min. Subsequently, DAPI working solution was applied and incubated at RT for 5 min in the dark. Finally, slides were washed four times with PBS for 5 min each and imaged using a Leica Mica confocal microscope with a 63x HC PL APO objective (Wetzlar, Germany) equipped with a Cy5 filter set.

### Measurement of cardiomyocyte respiration with Cell Mito Stress Test using Agilent Seahorse XF

Isolated cardiomyocytes from neonatal rats were seeded in 96-well plate at the density of 5000 cells/well for Seahorse assays (Agilent Seahorse XFE96 Analyzer). About 80% confluent cardiomyocytes were treated with IFNγ (10ng/ml) for 48h. Cardiomyocyte mitochondrial respiration was measured by determining oxygen consumption rate (OCR) using Agilent Seahorse XF Cell Mito Stress Test kit (Cat. #103015-100) following manufacturer guidelines.

### Immunofluorescence Staining

For immunofluorescence analysis, µ-slides were coated with collagen I prior to seeding NRVCMs or hiPSC-CMs. Following experimental treatments, cells were washed with PBS and fixed in 4% PFA (400 µL/well) for 10 min at 37 °C. After three PBS washes, cells were permeabilized with 0.1% Triton X-100 in PBS (400 µL/well) for 15 min at RT followed by three additional PBS washes.

Non-specific binding was blocked with 2.5% bovine serum albumin (BSA) in PBS (500 µL/well) overnight at 4 °C. Primary antibodies [rabbit anti-STAT1 polyclonal (1:500; #14994, Cell Signaling Technology, Taufkirchen, Germany), rabbit anti-GBP5 monoclonal (1:500; #313390, Abcam, Cambridge, UK), mouse anti-α-actinin monoclonal (1:5000; A7732, Merck KGaA, Darmstadt, Germany)] were diluted in 0.1% BSA/PBS and applied overnight at 4 °C. Slides were washed 3× with PBS and incubated with fluorophore-conjugated secondary antibodies [chicken anti-rabbit IgG Alexa Fluor 647, donkey anti-mouse IgG Alexa Fluor 546 (A10036; Thermo Fisher Scientific, Waltham, MA, USA)] in 0.1% BSA/PBS for 60 min at RT in the dark.

After three washes with PBS/0.05% Tween-20, nuclei were stained with DAPI (D9542, Merck KGaA; 1:5000 in PBS) for 5 min, followed by three final PBS washes. Images were acquired using a Leica MICA confocal microscope (63× HC PL APO objective, Wetzlar, Germany).

### Protein Preparation and Immunoblotting

To obtain protein from any used cell type in this project, RIPA lysis buffer (#9806 CST) together with protease and phosphatase inhibitor cocktail (Roche Applied Sciences, Penzberg, Germany), 0.1% SDS and 50 mM NaF was used. The cells were lysed by three freeze-thaw cycles in RIPA lysis buffer. To remove cell debris the lysates were centrifuged. Protein concentration was determined photometrically using a DC Assay kit (Bio-Rad, Feldkirchen, Germany). BSA was used as a standard. Protein lysates were denaturated using 4x Laemmli buffer at 95°C for 10 min. Protein samples were separated according to their molecular weight using 4-12% Criterion™ XT Bis-Tris Protein Gel (Bio-Rad, Feldkirchen, Germany), followed by transfer to nitrocellulose membrane and subsequent immunoblot with the respective target primary antibodies. The primary antibodies were incubated overnight at 4°C followed by incubation with a HRP-coupled secondary antibody (Santa Cruz Biotechnology, Dallas, TX, U.S.). Protein expression was visualized using an ECL based chemiluminescence kit (Cytiva Amersham™, Marlborough, Massachusetts, U.S.) and detected on a ChemiDoc MP Imaging System (Bio-Rad, Feldkirchen, Germany). The densitometric quantitative analysis was performed using ImageJ. All western blot experiments were performed in triplicates and repeated thrice.

### Antibodies

The antibodies used for various experiments in this study are the following: Caspase 7, rabbit polyclonal (Cat. no. 9492, 1:1000; Cell Signaling Technology, Germany); GBP5, rabbit monoclonal (Cat. No. EPR28367, 1:1000; Abcam; Cambridge, United Kingdom); STAT1, rabbit polyclonal (Cat. No. 14994, 1:1000; Cell Signaling Technology, Germany); p-STAT1, rabbit polyclonal (Cat. No. 9167, 1:1000; Cell Signaling Technology, Germany); Vinculin, rabbit polyclonal (Cat. No. 4650, 1:1000; Cell Signaling Technology, Germany), OXPHOS cocktail (Cat. No. ab110413, Abcam).

### Statistical analysis

Unless otherwise stated, all results shown are means ± SEM (Standard Error of the Mean). Data statistical analyses were conducted using a two-tailed Student’s t-test for each experimental analysis. Seahorse data was analysed with Unpaired t test with Welch correction. If required, a two-way ANOVA (followed by Student-Newman-Keuls post-hoc tests where appropriate) was performed. For all experiments, P value less than 0.05 was considered statistically significant.

## Acknowledgments

This research work was supported by DFG grants to M.K. (KU 4356/1-1). N.F. and T.S. acknowledge funding from DFG-SFB1550/1. We thank Rebecca Kistler for excellent technical support in providing hiPSC-derived cardiomyocytes. We thank Ashraf Rangrez and Anushka Deshpande for providing Gateway cloning vectors. We thank Marco Neu for help in conducting experiments for cardiomyocyte contractility assay. We thank Emelie Haas for technical support in TUNEL staining.

## Author Contributions

M.K. conceived the project and analyzed data. L.N. carried out all the experiments with help from L.La.. S.H. and T.S. provided hiPSC-derived cardiomyocytes and helped in conducting cardiomyocyte contractility assay. M.K. and L.N. analyzed data and wrote the manuscript. M.K. supervised all experiments. T.S., L.L., and N.F. discussed the results and edited the manuscript. N.F. T.S. and M.K. acquired funding.

## Supplementary information

The manuscript contains Supplementary Figures and a Table in supplemental data/

