## Supplemental files for "Cardiomyocytes execute pro- and anti-inflammatory signaling of IFNγ-induced GBP5 by differential regulation of the inflammasome"

**Figure S1. Cardiomyocytes stimulated with IFNγ upregulates GBP5.**

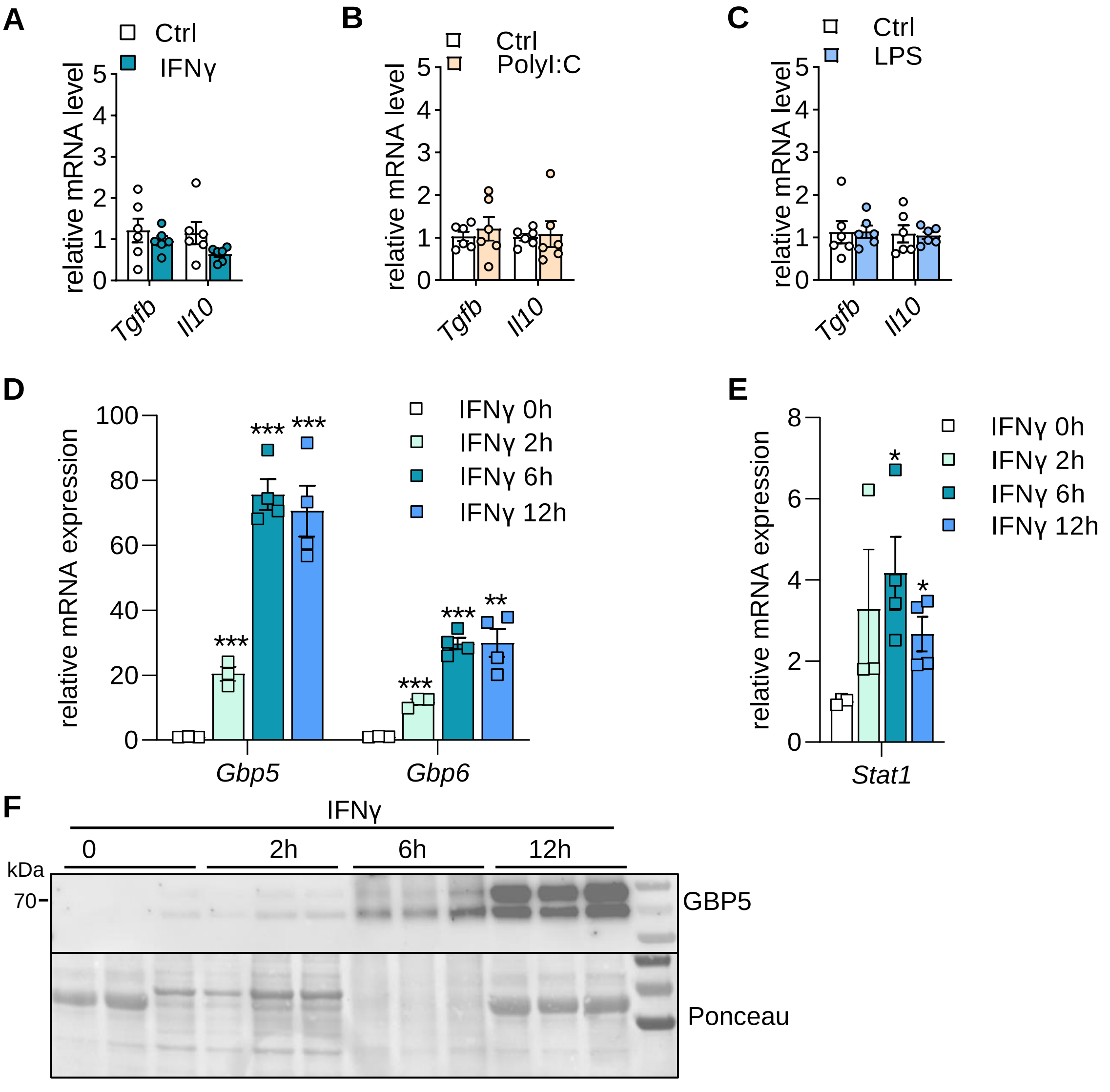


**Figure S1. Cardiomyocytes stimulated with IFNγ upregulates GBP5.**

**A-C**. Expression of *Tgfb* and *Il10* in NRVCMs upon type I (LPS and PolyI:C) and type II (IFNγ) IFN stimuli by qPCR. N=6/group.

**D-E**. Time dependent effect on *Gbp5*, *Gbp6*, and *Stat1* mRNA expression in NRVCMs upon IFNγ (10ng/ml) stimulation determined by qPCR. N=4/group. ***p<0.001, **p<0.01, *p<0.05.

**F**. Immunoblot showing time dependent effect on GBP5 protein levels in NRVCMs upon IFNγ (10ng/ml) stimulation. N=3/group.

**Figure S2. Differential effect of type I and type II IFN activation on hiPSC-Cardiomyocytes inflammasome signaling**

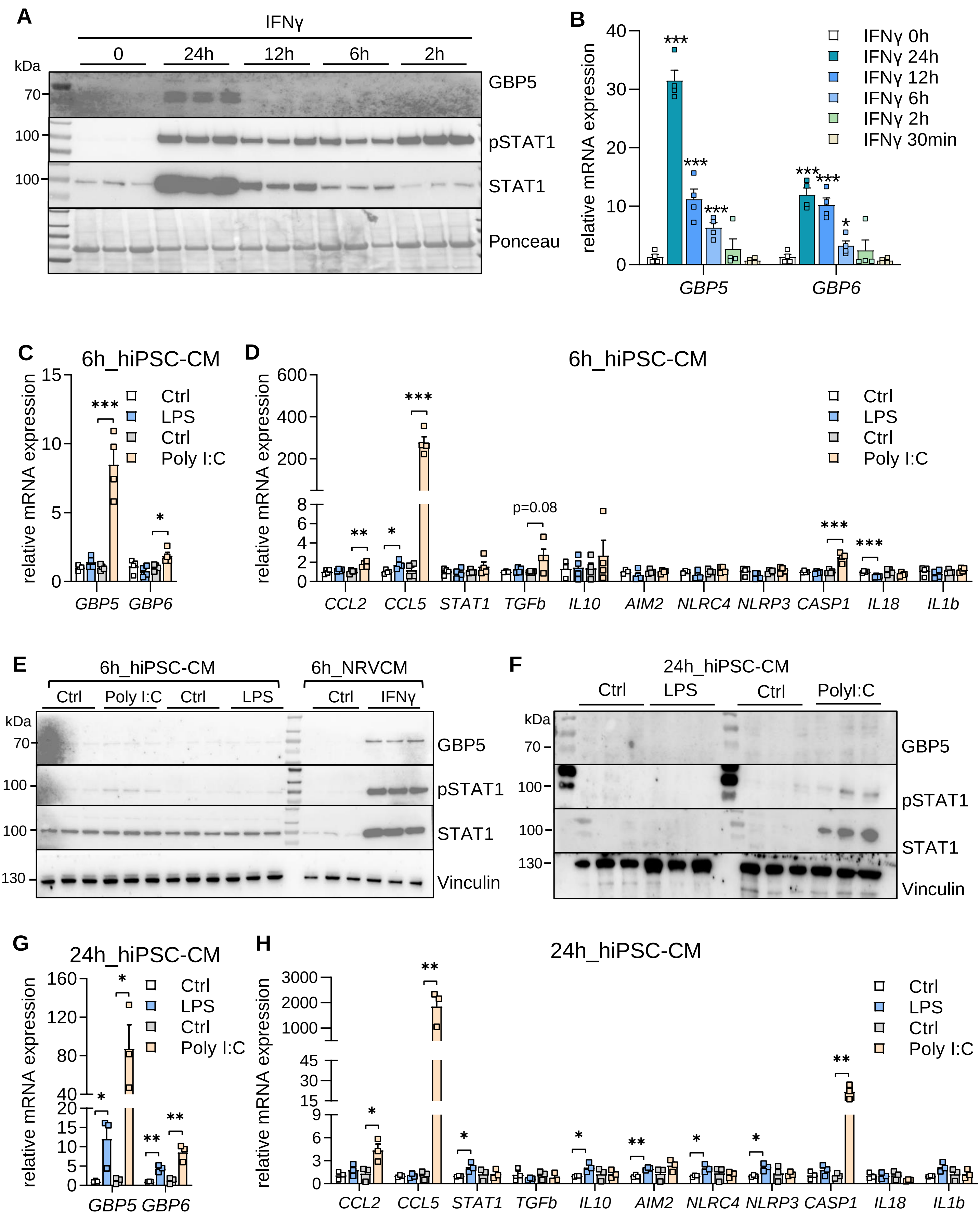

**Figure S2. Differential effect of type I and type II IFN activation on hiPSC-Cardiomyocytes inflammasome signaling.**

**A**. Immunoblot showing time dependent effect on GBP5 protein levels in hiPSC-CMs upon IFNγ (10ng/ml) stimulation. N=3/group.

**B**. Time dependent effect on *GBP5* and *GBP6* mRNA levels in hiPSC-CMs upon IFNγ (10ng/ml) stimulation determined by qPCR. N=4/group. ***p<0.001, *p<0.05.

**C**. Expression of GBP5 and GBP6 mRNA in hiPSC-CMs after 6h of LPS (100ng/ml) and poly I:C (1μg/ml) stimulation. N=4/group. ***p<0.001, *p<0.05.

**D**. Effect on inflammatory marker genes in hiPSC-CMs after 6h of LPS (100ng/ml) and poly I:C (1μg/ml) stimulation. N=4/group. ***p<0.001, **p<0.01, *p<0.05.

**E-F**. Immunoblot showing GBP5, pSTAT1, and STAT1 protein levels in hiPSC-CMs after 6h and 24h treatment with LPS (100ng/ml) and poly I:C (1μg/ml).

**G**. Expression of GBP5 and GBP6 mRNA in hiPSC-CMs after 24h of LPS (100ng/ml) and poly I:C (1μg/ml) stimulation. N=3/group. ***p<0.001, *p<0.05.

**H**. Effect on inflammatory marker genes in hiPSC-CMs after 24h of LPS (100ng/ml) and poly I:C (1μg/ml) stimulation. N=3/group. ***p<0.001, *p<0.05.

### Figure S3. Effect of IFNγ on cardiomyocyte cell contractility.

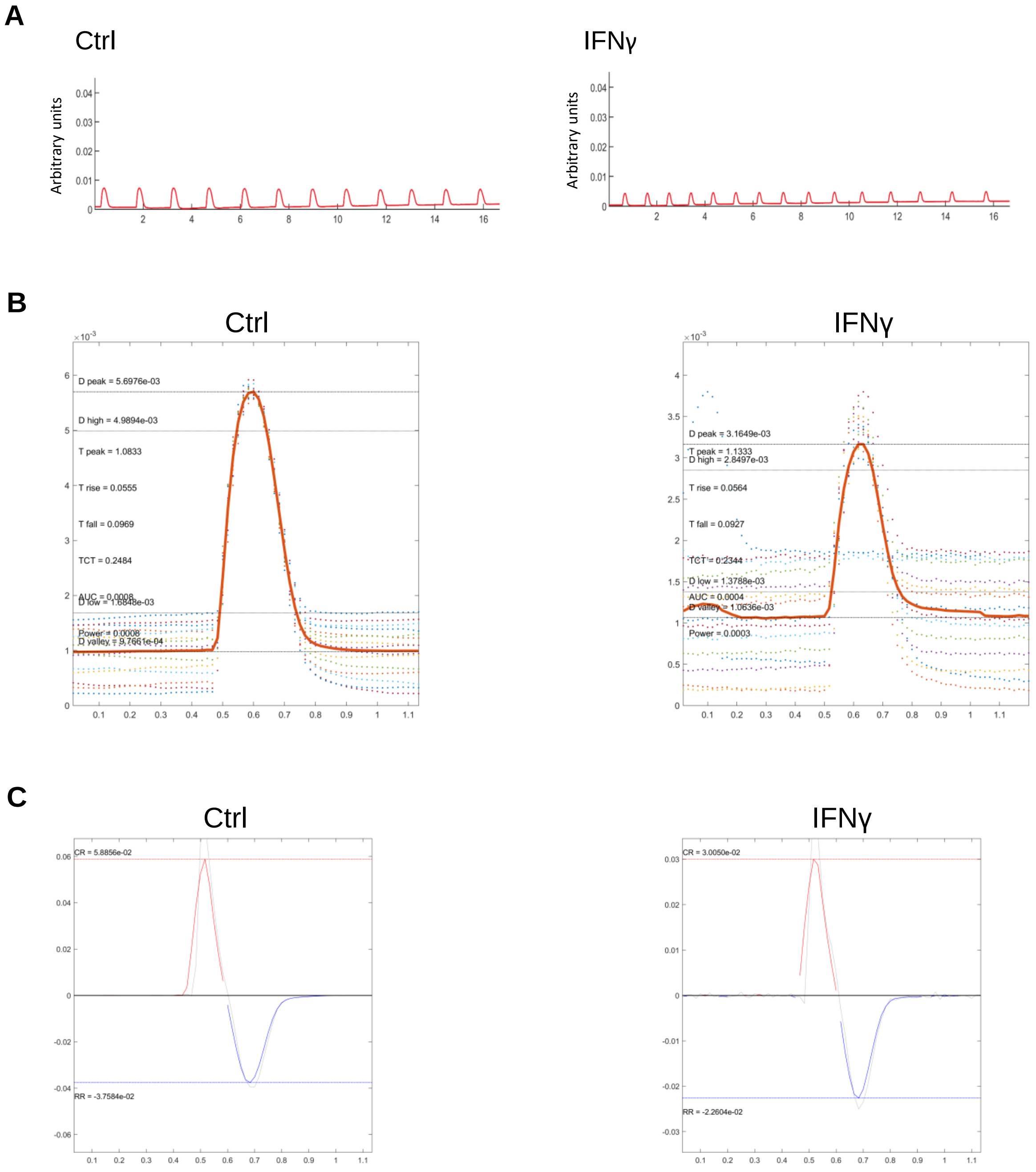


**Figure S3. Effect of IFNγ on cardiomyocyte contractility.**

**A-C.** Cardiomyocyte from neonatal rats were treated with varying concentrations of IFNγ (10ng/ml) for 48 h and contractility was assessed as described in the method section. Representative pictograms showing T_Peak (time to peak contraction), D_Peak (duration of peak contraction), contraction velocity (rate of cell shortening), and relaxation velocity (rate of cell lengthening) in cardiomyocytes are shown.

**Figure S4. Effect of IFNγ concentration on cardiomyocyte cell viability.**

**A**

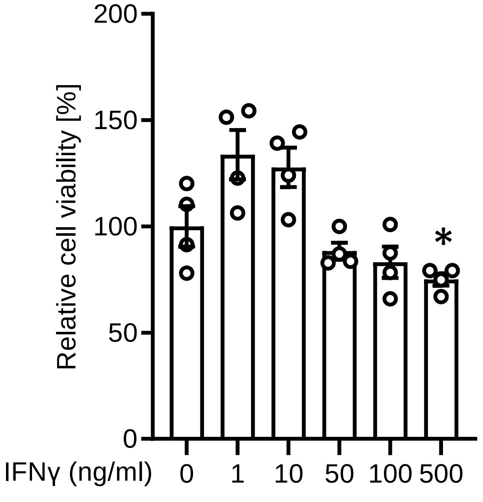


**Figure S4. Effect of IFNγ concentration on cardiomyocyte cell viability.**
**A**. Cardiomyocyte were treated with varying concentrations of IFNγ for 48 h and cell viability was assessed by MTT assay kit following manufacture's instructions. N=4/group. *p<0.05.
